# Genetic heterogeneity alleviates behavioral trade-off between exploration and vigilance in *Drosophila*

**DOI:** 10.1101/2025.10.16.682765

**Authors:** Takahira Okuyama, Daiki X. Sato, Yuma Takahashi

## Abstract

Trade-offs between conflicting behaviors fundamentally constrain animal performance, yet group living can alter how such trade-offs are expressed and alleviated. Here, we disentangled the effects of aggregation and heterogeneity, both arising from group living, on the exploration–vigilance trade-off in *Drosophila*. Using 78 genetically distinct strains from the *Drosophila* Genetic Reference Panel, we compared behavioral responses of single individuals, homogeneous groups (six individuals from the same strain), and heterogeneous groups (three individuals from each of two distinct strains) under predator-like looming stimuli. Pareto optimization analysis revealed that aggregation increased both exploratory and vigilance behavior, thereby exposing the underlying exploration–vigilance trade-off. In contrast, heterogeneity enhanced group performance beyond expected levels, alleviating the cost of balancing exploration and vigilance. To identify the phenotypic basis of this alleviation, we developed a phenome-wide higher-level association study (PheHAS) that screened 190 phenotypes across strain pairs. We found that heterogeneity in activity patterns and morphological traits promoted trade-off alleviation, whereas similarity in infection responses also contributed, suggesting that social facilitation may contribute. Our study provides the first explicit demonstration that aggregation and heterogeneity exert distinct effects on a group-level behavioral trade-off and introduces PheHAS as a generalizable framework for identifying the phenotypic bases underlying diversity effects in animal groups.

## Introduction

Life-history traits are constrained by many trade-offs that arise when one trait cannot increase without a decrease in another (1, 2). This concept is crucial in understanding how animals balance competing demands for multiple resources throughout their life. Prior studies have categorized trade-offs into two types, so-called “allocation trade-off” and “acquisition trade-off” (3). Allocation trade-offs typically arise when animals distribute their limited resources (4, 5) and lead to a negative correlation between life history traits such as reproduction and survival (6, 7). However, many empirical studies have reported positive or negligible correlations between these traits, rather than negative correlations predicted by theory of allocation trade-off (8, 9). A long-term monitoring study demonstrated that female reindeer with higher reproductive success also exhibited higher survival rates (10). These studies highlight the importance of the concept of “acquisition”, meaning that individuals acquiring more resources can allocate more energy to multiple traits. However, some studies have shown that acquisition often comes at a cost (11, 12). While animals strive to maximize their resource utilization, phenotypic correlations can sometimes constrain them from reaching the optimum, resulting in negative correlations between traits. This phenomenon, known as the acquisition trade-off, has been an important topic in behavioral ecology (13).

The foraging–vigilance trade-off is one of the most typical examples of acquisition trade-offs, and extensive field and experimental studies have demonstrated its existence and underlying mechanisms (14, 15). Organisms face potential risks of both starvation and predation. Since foraging and avoiding predators cannot be performed simultaneously, a conflict arises in deciding how much to prioritize one behavior over the other. For example, a study using goldfish (*Carrassius auratus*) as preys and little egrets (*Egretta garzetta*) as predators showed that individuals more motivated to forage were more likely to be preyed upon (16). However, unlike allocation trade-offs, acquisition trade-offs can sometimes be alleviated through cooperation in group-living animals. For instance, ostriches cannot maintain vigilance while foraging with their heads down, facing an acquisition trade-off between vigilance and foraging in solitary conditions. In contrast, within a group, individuals can focus on foraging while others remain vigilant (17). This phenomenon, where individuals increase time spent on foraging without decreasing time spent on vigilance as group size increases, has been observed in some species (18, 19). In other words, living in groups has the potential to resolve the foraging–vigilance trade-off.

In addition to the effects of aggregation, grouping may also exert the effects of heterogeneity. Accumulating evidence suggests that interindividual heterogeneity in behavior can drive emergent changes in group performance (20). For example, groups of mice foraged more efficiently than individuals due to the presence of key individuals that were particularly good at foraging (21). Similarly, fly groups comprised of two distinct strains exhibited enhanced exploration, likely due to differences in locomotor activity levels between genotypes (22). Such phenomena are partially explained by the well-established concept of indirect genetic effects, whereby the phenotype of group members influence an individual’s phenotype (23, 24). While these findings highlight the significant role of heterogeneity in group dynamics, no study has simultaneously examined multiple behavioral phenotypes or assessed how heterogeneity influences their trade-off relationships. Furthermore, the effects of aggregation and heterogeneity have not been explicitly distinguished or directly compared in terms of how group composition influences animal behavior. Integrating the impacts of aggregation and heterogeneity offers a clearer understanding of how groups navigate behavioral trade-offs in nature.

Here, we investigate how group composition shapes the behavioral trade-off between exploration and vigilance using 78 genetically distinct strains of *Drosophila melanogaster* derived from the *Drosophila* Genetic Reference Panel (DGRP) (25). We set up three social conditions: single individuals, homogeneous groups (composed of a single strain) and heterogeneous groups (composed of two distinct strains). Behavioral recordings were conducted for 5 minutes under conditions with and without looming stimuli. We quantified exploratory behavior as search comprehensiveness and vigilance behavior as freezing rate. Pareto optimization analysis was applied to quantify and compare the relative performance of flies across the social conditions under the behavioral trade-off. Furthermore, we identified the phenotypic diversity potentially resolving the trade-off using a comprehensive database. Our study provides novel insights into how group composition enhances behavioral performance by alleviating the trade-off between exploration and vigilance.

## Results

### Group composition alters mean levels of exploration and vigilance

To assess how group composition affects individual behavior, we compared the mean values of exploration and vigilance traits across three social conditions. For exploratory behavior, a significant difference was observed between single individuals and homogeneous groups (*W* = 247, *df* = 78, *P* < 0.001), while no significant difference was found between expected values— calculated as the mean of the corresponding homogeneous groups—and observed values in heterogeneous groups (*W* = 1096, *df* = 65, *P* = 0.88), suggesting that aggregation promotes resource exploration (Fig. 2A).

**Figure 1.**
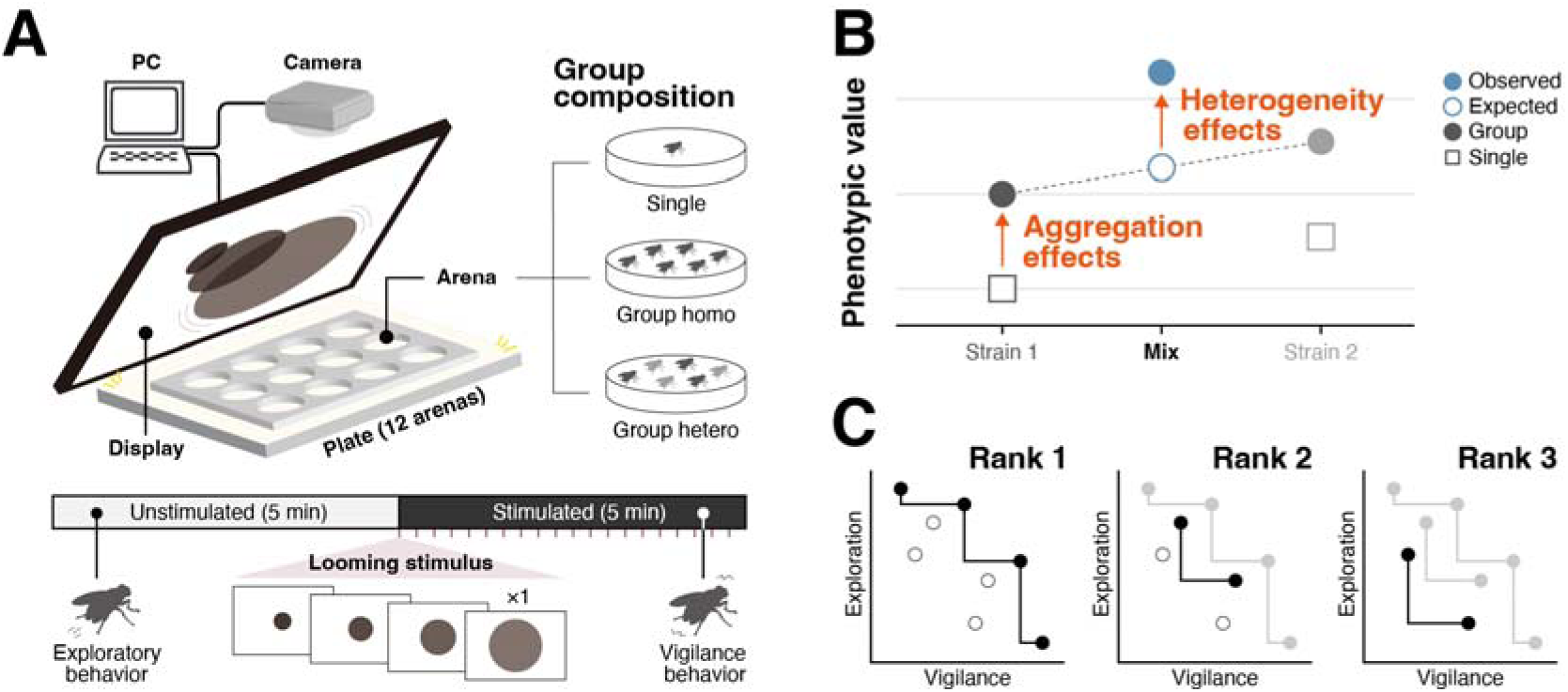
Experimental scheme and analytical framework. **(A)** Flies were tested in experimental arenas placed under a display presenting looming stimuli. Behavioral data were collected under three social conditions: single individuals, homogeneous groups of six from the same strain, and heterogeneous groups of three each from two strains. Unstimulated periods (5 min) were used to measure exploratory behavior, and stimulated periods (5 min) to measure vigilance. **(B)** Calculation of aggregation and heterogeneity effects. Aggregation effects were the deviation of group phenotypic values from the mean of single individuals, whereas heterogeneity effects were the deviation of observed heterogeneous groups from their expected values based on homogeneous groups. **(C)** Pareto optimization analysis for evaluating the exploration–vigilance trade-off. Strategies located in the upper-right region of the exploration–vigilance space (i.e., high exploration and vigilance) were assigned rank 1, with subsequent ranks determined iteratively. This provides a quantitative measure of how group composition influenced trade-off resolution.

**Figure 2.**
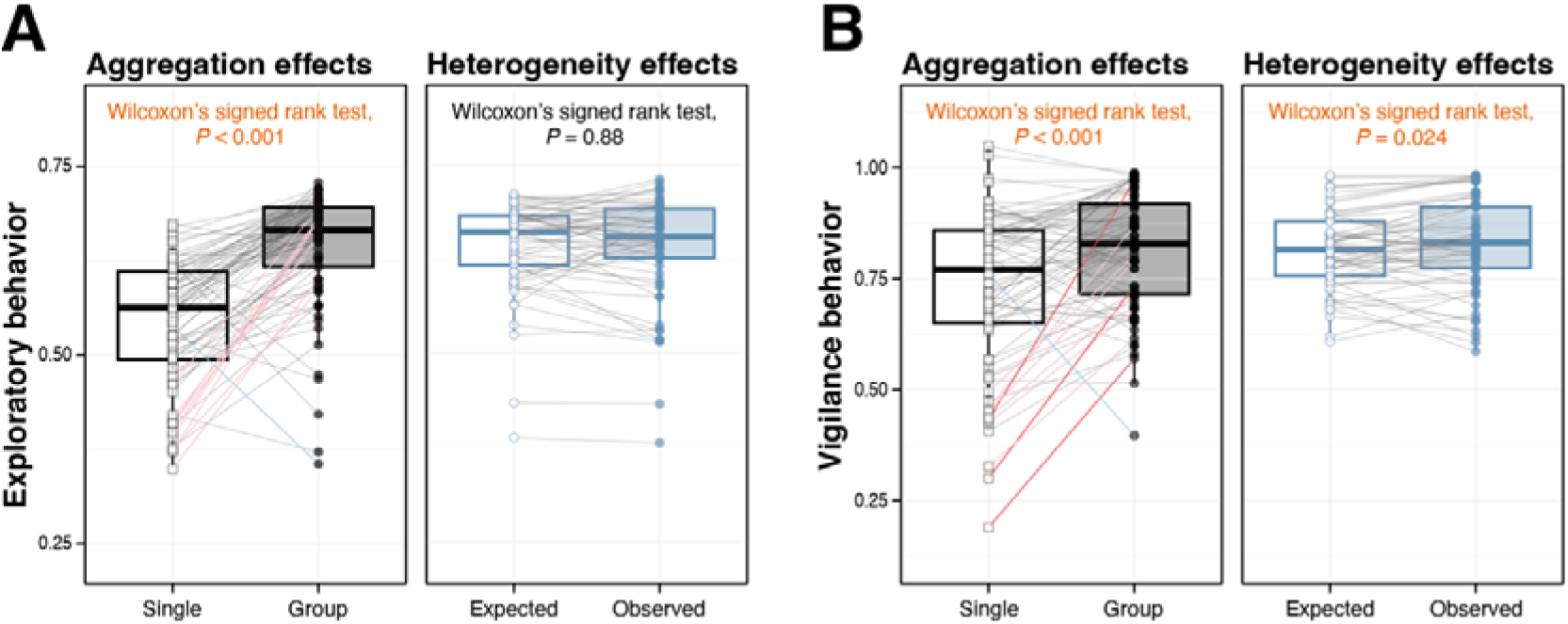
Aggregation and heterogeneity effects on exploratory and vigilance behavior. **(A)** In exploratory behavior, homogeneous groups showed significantly higher values than single individuals, indicating an aggregation effect. No significant difference was observed between expected and observed values in heterogeneous groups. **(B)** In vigilance behavior, both aggregation and heterogeneity effects were detected: homogeneous groups differed from single individuals, and observed heterogeneous groups exceeded their expected values. Although overall effects tended to be positive, their magnitude varied considerably across strains and strain pairs (see Fig. S1). Lines connect corresponding strains or strain pairs across conditions, with colors indicating fold change: red, >2-fold increase; pink, 1.5–2-fold increase; blue, >2-fold decrease; light blue, 1.5–2-fold decrease; gray, all other cases.

In contrast, for vigilance behavior, significant differences were observed both between single individuals and homogeneous groups (*W* = 687, *df* = 78, *P* < 0.001) and between expected and observed values in heterogeneous groups (*W* = 727, *df* = 65, *P* = 0.024), indicating that both aggregation and heterogeneity may contribute to anti-predator response (Fig. 2B). Although the overall direction of these effects were generally positive, their magnitude varied substantially across strains and strain pairs (Fig. S1).

### Effects of aggregation and heterogeneity on the exploration–vigilance trade-off

To examine the effect of group compositions on the relationship between exploration and vigilance, we compared the correlation between these traits across social conditions. Under single-individual conditions, no significant correlation was observed (*r* = −0.091, *P* = 0.43), whereas under homogeneous group conditions, a significant negative correlation emerged (*r* = −0.48, *P* < 0.001), indicating the presence of exploration–vigilance trade-off. To reveal the effects of group composition on the behavioral trade-off, we applied Pareto optimization analysis and obtained Pareto ranks of each strain (Fig. 3A). Pareto ranks were significantly lower under homogeneous group conditions than under single-individual conditions (*W* = 2850, *df* = 78, *P* < 0.001), indicating higher performance conferred by aggregation (Fig. 3B). Under heterogeneous group conditions, even stronger negative correlations between vigilance and exploration were detected for both expected (*r* = −0.49, *P* < 0.001) and observed values (*r* = −0.50, *P* < 0.001) (Fig. 3C), confirming the presence of exploration–vigilance trade-off. Pareto ranks from observed values were significantly lower than those from expected values (*W* = 1435, *df* = 65, *P* < 0.001), indicating that heterogeneity within group substantially enhanced behavioral performance under the trade-off (Fig. 3D).

**Figure 3.**
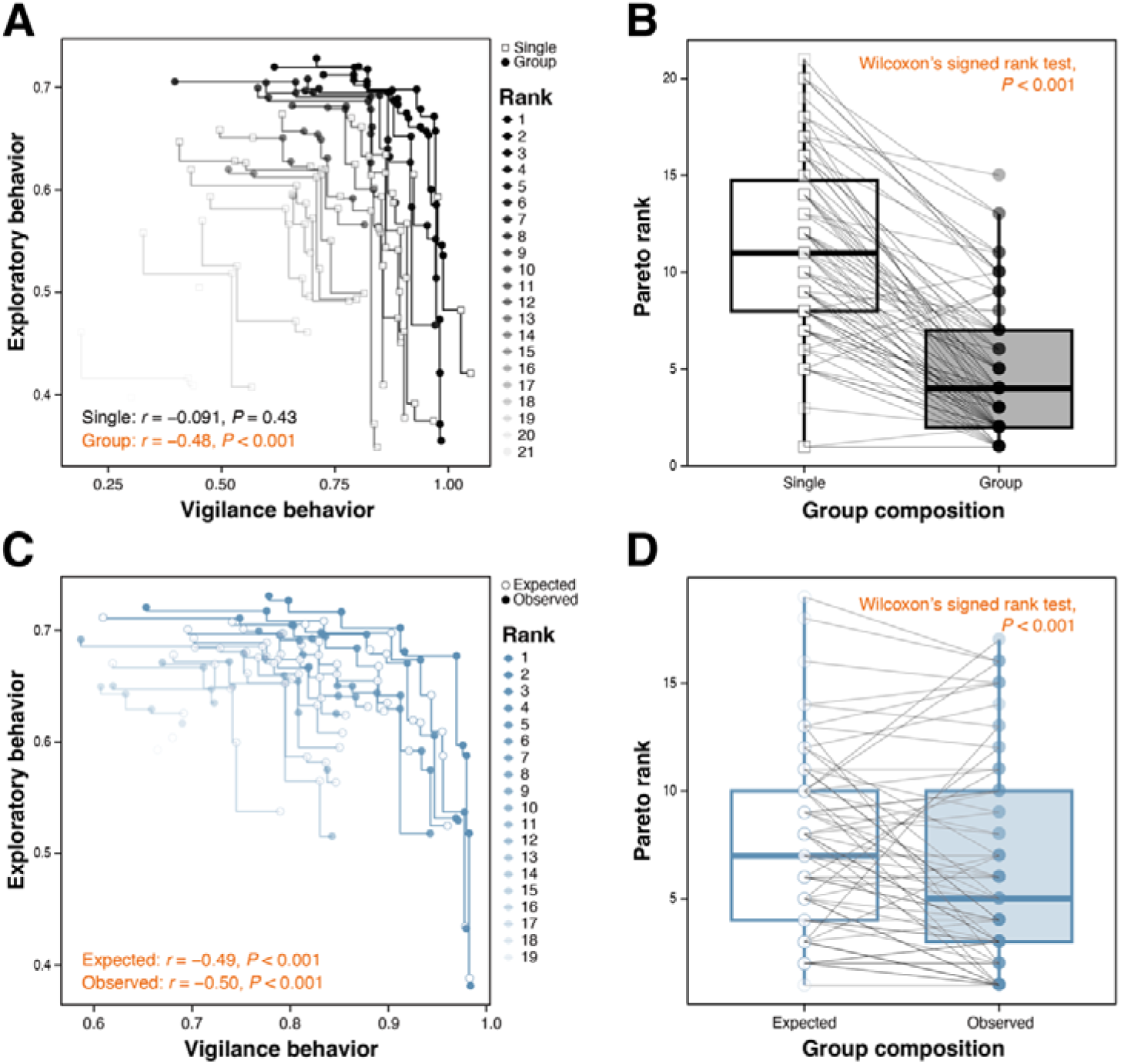
Effects of aggregation and heterogeneity on the exploration–vigilance trade-off. **(A)** Relationship between exploration and vigilance under single-individual and homogeneous group conditions. A clear negative correlation emerged in groups, indicating an exploration–vigilance trade-off. **(B)** Pareto optimization analysis showed that homogeneous groups achieved higher ranks than single individuals. **(C)** Relationships between exploration and vigilance in expected and observed heterogeneous groups, both showing significant negative correlations. **(D)** Pareto ranks were higher in observed than expected heterogenous groups, indicating trade-off alleviation driven by heterogeneity effects.

### Phenotypic heterogeneity underlies alleviation of the behavioral trade-off

To examine how phenotypic variation among individuals relates to the degree of alleviation of the exploration**–**vigilance trade-off, we conducted a phenome-wide association analysis using 190 phenotypes of the 73 DGRP strains used in this study, obtained from the DGRPool database (26). Hierarchical clustering grouped the 190 phenotypes into 16 clusters, which were further classified into four higher-order categories: physiology, morphology, cuticular component, and life history (Fig. S2). Analogous to the genome-wide higher-level association study (27), which examines associations between the genetic characteristics of a group and its higher-level traits (e.g., diversity effects arising from the genetic diversity of its members), we refer to the present analysis as a phenome-wide higher-level association study (PheHAS).

Pairwise distances between strains were calculated for each phenotype, and we examined their correlations with the trade-off alleviation scores. To assess whether such associations were consistent within related traits, correlation coefficients were analyzed at the clusters level (Fig. 4). In the physiological category, the activity pattern cluster showed a significant positive correlation overall (*r* = 0.18, *P* < 0.001), indicating that greater differences in activity patterns were associated with stronger alleviation of the exploration–vigilance trade-off. In contrast, the infection response cluster exhibited a significant negative correlation (*r* = −0.16, *P* = 0.0049), suggesting that smaller differences in infection responses contributed to better trade-off alleviation. The startle response cluster showed no significant correlation (*r* = 0.016, *P* = 0.75). At the phenotype level, waking activity (*r* = 0.27, *P* = 0.046) and sleep bout duration (*r* = 0.40, *P* = 0.0030) displayed particularly strong positive correlations.

**Figure 4.**
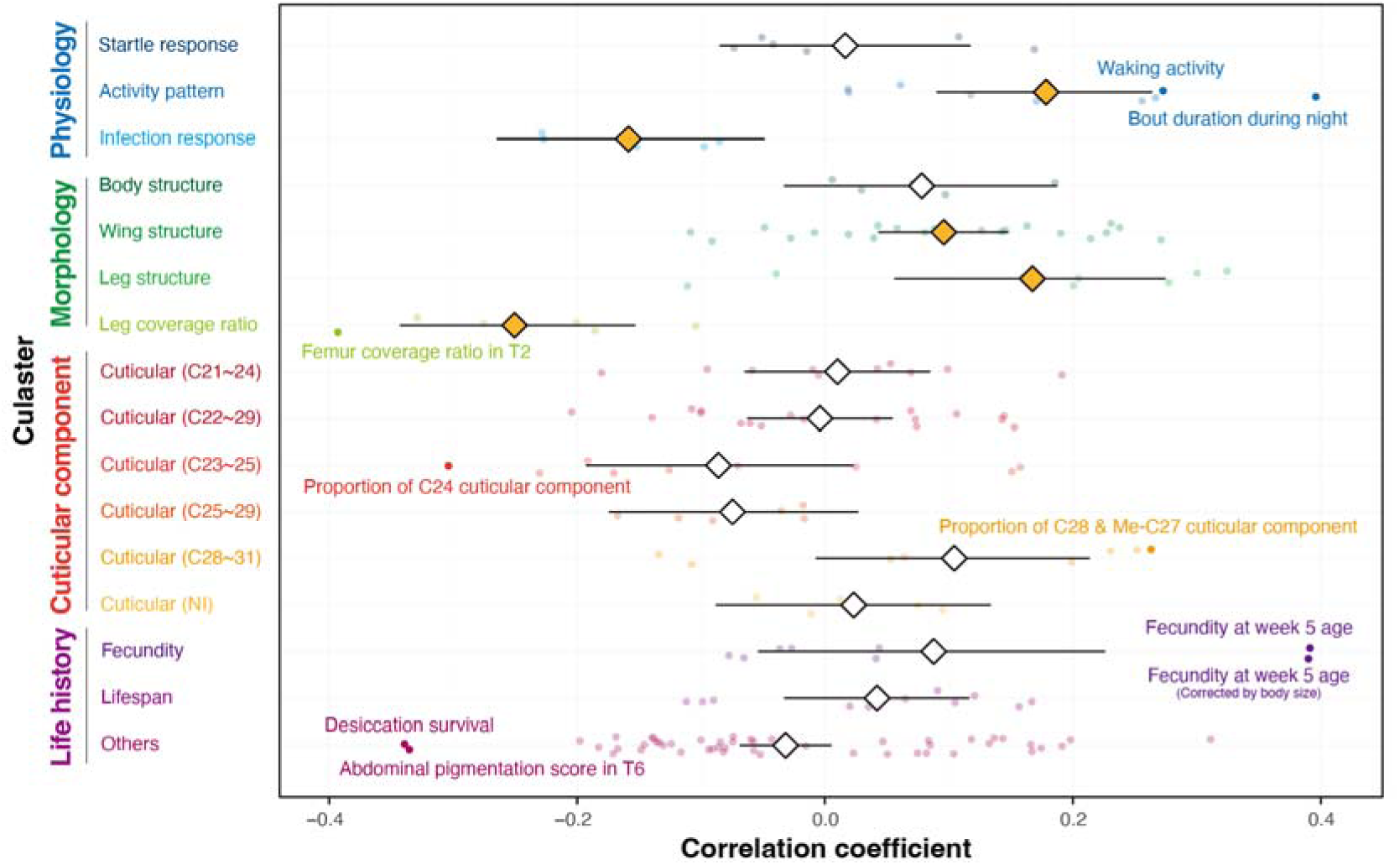
Phenotypic bases of trade-off alleviation identified by Phenom-wide higher-level association study (PheHAS). Correlation coefficients between phenotypic distances and trade-off alleviation scores were examined across 190 traits, grouped into 16 clusters and four higher-order categories: physiology, morphology, cuticular components, and life history. Differences in several traits and clusters were significantly associated with trade-off alleviation, highlighting the phenotypic bases through which group composition reshapes behavioral trade-offs.

In the morphological category, clusters related to wing (*r* = 0.096, *P* < 0.001) and leg structure (*r* = 0.17, *P* = 0.0034) showed significant positive correlations, indicating that greater differences in these structures were associated with alleviation of the exploration–vigilance trade-off. In contrast, the leg coverage ratio cluster exhibited a significant negative correlation (*r* = −0.25, *P* < 0.001), suggesting that more similar coverage ratios promoted better trade-off alleviation. No significant correlation was detected for the body structure cluster (*r* = 0.078, *P* = 0.17). At the phenotype level, the femur coverage ratio in the second thoracic segment (T2) showed a strong negative correlation (*r* = −0.39, *P* = 0.032).

In the cuticular component category, the C28–C31 cluster showed a weak positive correlation (*r* = 0.10, *P* = 0.068), indicating that greater variation in the proportion of these components contributed to alleviation of the exploration–vigilance trade-off. Other cuticular clusters (C21–C24, C22–C29, C23–C25, C25–C29, and NI) showed no significant correlations. At the phenotype level, the proportions of C28 and Me-C27 components exhibited significant positive correlations (*r* = 0.26, *P* = 0.048), whereas the proportion of C24 showed a significant negative correlation (*r* = −0.30, *P* = 0.022).

In the life history category, no significant correlations were observed at the cluster level, including fecundity (*r* = 0.088, *P* = 0.23), lifespan (*r* = 0.042, *P* = 0.27) and other life-history traits (*r* = −0.032, *P* = 0.094). At the phenotype level, fecundity at 5 weeks (*r* = 0.39, *P* = 0.0039) and its corrected value (*r* = 0.39, *P* = 0.0038) were positively correlated with alleviation score, whereas desiccation survival (*r* = −0.34, *P* = 0.0080) and abdominal pigmentation score in T6 (*r* = −0.34, *P* = 0.0094) were negatively correlated.

## Discussion

Aggregation and heterogeneity jointly shape group behavior. Aggregation effects are well established through density-dependent mechanisms such as the dilution effect (28) and confusion effect (29), where group size alters risk and decision-making. Heterogeneity effects, in contrast, arise from qualitative differences among individuals, such as indirect genetic effects (23, 24), in which partner traits influence behavioral outcomes. However, most prior work has faced two limitations: conflating aggregation with heterogeneity, and examining individual behaviors in isolation. These limitations obscure trade-off structures and behavioral syndromes. Using *D*. *melanogaster*, we disentangled these effects and revealed their phenotypic basis.

Homogeneous groups showed greater spatial coverage than single-individual conditions, revealing a clear aggregation effect in exploratory behavior. Enhanced coverage of exploration likely increases encounters with food, mates, or oviposition sites. Such improvements associated with group living may stem from social facilitation, in which the presence of conspecifics enhances the intensity of certain behaviors (30, 31). In *Drosophila*, social facilitation has also been reported in contexts such as grooming (32) and long-term memory (33, 34), suggesting that our observations may represent another manifestation of this phenomenon. A parallel aggregation effect was observed in vigilance: homogeneous groups froze more frequently than individuals, indicating enhanced anti-predator responses through social interaction (27). Freezing is a widespread defensive strategy observed across mammals (35), fish (36), and insects (37), and often scales with group size. The many-eyes hypothesis (38) proposes that larger groups detect predators earlier by distributing vigilance, while the Trafalgar effect (39) highlights the role of social cues in early detection of threats not directly perceived by all individuals. Taken together, these mechanisms provide a compelling explanation for the enhanced vigilance observed in our study. Beyond aggregation, heterogeneity also shaped vigilance. Heterogeneous groups froze more than expected from homogeneous groups, revealing a heterogeneity effect, consistent with our prior findings (27). In vertebrates, sentinel systems emerge when certain individuals monitor while others forage, thereby boosting group vigilance (40, 41). Although such structured systems are unexplored in flies, variation in sensitivity thresholds could similarly allow some individuals to benefit from the vigilance of more reactive others.

Group composition also reshaped behavioral trade-offs. No correlation between exploration and vigilance appeared in the single-individual condition, but a strong negative correlation emerged in homogeneous groups. Garland (2014) illustrated this phenomenon with speed and stamina: trade-offs may remain hidden among average performers but become evident among top athletes (2). This suggests that the visibility of trade-offs depends on the group’s overall performance level—higher performance can make hidden trade-offs detectable. By analogy, aggregation raised both exploration and vigilance, likely driven by social facilitation and the many-eyes effect, thereby revealing the underlying trade-off. Pareto optimization analysis confirmed higher performance under group conditions compared to individuals, but aggregation alone did not resolve the trade-off. In heterogeneous groups, observed performance exceeded expected performance despite the persistence of the trade-off, demonstrating that heterogeneity can alleviate the cost of balancing exploration and vigilance. While trade-off alleviation has been discussed in ecology (42) and evolution (43), it has not been explicitly demonstrated in animal behavior. Our findings therefore provide novel insights into how genetic heterogeneity can help resolve fundamental dilemmas animals face in their lives.

Using our novel framework, PheHAS, we identified within-group phenotypic diversity that alleviates the exploration–vigilance trade-off. In the physiological category, greater heterogeneity in activity patterns predicted stronger alleviation of the trade-off. The response threshold model (44, 45) explains how inter-individual variation in task-specific thresholds can drive division of labor; extensions including copying behavior (46, 47) and observational conditioning (48, 49) suggest that diversity in sensitivity enables flexible responses to environmental cues. From this perspective, relatively insensitive individuals—often described as bold—may promote exploration in their groupmates, whereas more sensitive individuals—often described as shy—may induce freezing in others, allowing the group as a whole to balance exploration and vigilance. Conversely, lower heterogeneity in infection responses correlated with greater trade-off alleviation. In social insects, cooperation and synchronized collective behavior enhance resistance to parasites and pathogens, a phenomenon known as social immunity (50, 51). Reduced heterogeneity in infection responses may therefore reflect stronger social cohesion, potentially reinforcing social facilitation (33, 30). This amplification may indirectly contribute to resolving the trade-off, although this interpretation warrants caution given the small sample size of this cluster.

Beyond physiology, several other traits also contributed to trade-off alleviation. Morphological traits (wing structure, leg structure, and leg coverage ratio), whose heterogeneity was positively correlated with alleviation, may influence locomotion (flying (52); walking (53)) and social behaviors (aggression (54); grooming (55)). Cuticular components, important in hormonal regulation and kin recognition (56, 57), showed a weak positive correlation only in the structurally complex C28–31 cluster. These results suggest that the effects of heterogeneity may depend on the ability of individuals to recognize whether conspecifics are genetically related. In life history traits, fecundity and desiccation survival were also correlated with alleviation, indicating that both short-term behavioral or physiological and long-term life-history traits shape group performance. Nevertheless, the specific interactions underlying these correlations remain hypothetical.

Altogether, we show that aggregation and heterogeneity exert distinct, measurable effects on group behavior and identify their phenotypic bases. To our knowledge, this is the first study to disentangle aggregation from diversity effects and to demonstrate that genetic heterogeneity can alleviate a fundamental behavioral trade-off between exploration and vigilance. PheHAS provides a generalizable framework for uncovering the traits through which such alleviation occurs. These findings suggest that trade-offs, rather than being immutable constraints, can sometimes be alleviated—a perspective that may help explain how evolution overcomes behavioral dilemmas. Genetic diversity may represent one such evolutionary strategy. Elucidating the functions and mechanisms of heterogeneity effects will offer deeper insights into the dynamics of animal group behavior.

## Materials and Methods

### Fly strains and rearing conditions

We used 78 inbred strains of *Drosophila melanogaster*, all derived from the *Drosophila* Genetic Reference Panel (DGRP), which was established from a natural population in the United States (25). Flies were maintained under standardized rearing conditions (12L:12D cycle, 25°C, ∼50% humidity) on standard cornmeal food (Bloomington Formulation, Nutri-Fly™, #66-113, Genesee Scientific), which also served as the substrate for egg laying and larval development.

To avoid sexual behavior that could interfere with responses to conspecifics and predator-like stimuli, all experiments used single-sex groups. Flies were collected 0–2 days after eclosion and reared for an additional 2 days under the same conditions. During this period, 5 males and 15 females were housed together in a single vial to ensure mating and to randomize re-mating experience. This was based on previous reports indicating that most flies are mated in natural environments (58) and that mating history can significantly influence behavioral patterns (59, 60). Adult females aged 2–4 days were used for the behavioral assay.

### Behavioral assay and tracking

Behavioral assays were conducted primarily during Zeitgeber time 6–12, corresponding to the daytime portion of the flies’ light-dark cycle (lights on at ZT 0, lights off at ZT 12). The incubator conditions were kept consistent with rearing (12L:12D, 25°C, ∼50% humidity). Prior to the experiment, females were briefly anesthetized with CO_2_ and introduced into the experimental arenas, which were 30 mm in diameter and 2 mm in depth, sufficient to prevent flight.

The experimental setup followed previous methods (61), as illustrated in Fig. 1A. After an initial 30-min recovery period from anesthesia, 12 arenas arranged on a single acrylic plate were placed onto an LED board (147 mm × 115 mm, 7500 lx white light; TLB-MP, Asone Co., Japan) and given an additional 10 min for acclimation. Subsequently, fly behavior was recorded for 10 min.

During the last 5 min of recording, flies were exposed to looming stimuli consisting 20 trials of a black circle expanding over 500 ms on a white background, presented every 15 s, following previous protocols (62, 63, 61). The stimuli were displayed at 60 Hz on a 13.3-inch EVICIV monitor, tilted at a 45° angle toward the arenas.

Locomotive behavior of flies was recorded at 640 × 480 pixel resolution and ∼50 frames per second using a USB3 camera (DMK33UX290, The Imaging Source, Germany) controlled by a custom Python script. The recorded videos were cropped to isolate each arena, and fly trajectories were extracted using Flytracker (64).

### Indices of exploration and vigilance

Tracked coordinates were standardized based on the position of the arena, such that the center of the arena was set to (0,0) and the diameter to 30. From the standardized coordinates, the distance traveled from the previous time point and the speed (distance divided by frame interval) were calculated at each time point and summarized into 0.5-s bins.

We then quantified both exploration and vigilance. The exploration index was measured as the evenness of spatial coverage during an unstimulated 5-minute period, as previously described (22). To calculate this, the arena was divided into five concentric regions of equal area, and Simpson’s diversity index (65) was computed based on the time spent in each region.

The vigilance index was measured as the rate of freezing in response to predator-like looming stimuli during a 5-min stimulation period. Freezing was defined as a reduction of movement speed below 4 mm/s within 0.5 seconds of stimulus onset, and the freezing rate was calculated as the proportion of trials in which this criterion was met.

### Calculation of the aggregation effect and heterogeneity effect

To examine how flies’ behavior changes with aggregation and heterogeneity among individuals, we compared their behaviors across three social conditions (Fig. 1B). This analysis highlighted the aggregation and heterogeneity effects, i.e., the synergistic effects of interindividual interaction on group performance. We collected behavioral data from 78 inbred strains of *D. melanogaster*, including both single individuals and homogeneous groups (6 individuals of the same strain per group). In addition, we analyzed 65 heterogeneous groups, each composed of two distinct strains, with 3 individuals from each strain.

The aggregation effect for a given strain was calculated as the discrepancy between phenotypic values of single individuals and homogeneous groups, as follows:

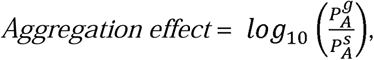

Where 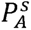 represents the mean phenotypic value of single individuals, and 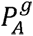 represents that of the homogeneous group.

Referring to a previous study (22), the heterogeneity effect for a given mixed-strain group was calculated as follows:

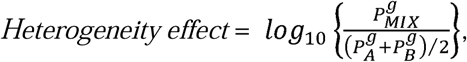

where 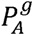 and 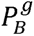 represent the mean phenotypic values of the homogeneous groups of the two strains comprising the heterogeneous group, and 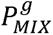 represents the phenotypic value of the heterogeneous group.

To calculate the mean phenotypic values for each social condition (i.e., 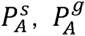, 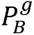, and 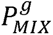), we used a linear mixed model with behavioral indices as the response variable, strain identity as a fixed effect, and arena position, plate identity, and time of day as random effects. Experiments for each strain or strain pair were conducted 17-20 times for single individuals and 3–6 times for groups, with the arena position within the plate randomized for each repetition.

### Evaluating exploration and vigilance trade-off

We first defined “behavioral strategy” of a given strain and/or strain pair based on its coordinate on the two-dimensional plane of exploration and vigilance. We then applied Pareto optimization analysis, a method widely used in engineering and economics (66), to evaluate the relative performance of it. Specifically, as illustrated in Fig. 1C, strain pairs that had no other strategies located in the upper-right region—representing simultaneously higher exploration and vigilance—were assigned to rank 1. These rank 1 strategies were then removed, and the same procedure was repeated to assign rank 2, rank 3, and so on. This ranking approach allowed us to identify strategies that resolved the trade-off between exploration and vigilance and thus were assigned higher ranks. Using these ranks, we evaluated how group composition influenced the structure of this behavioral trade-off.

### Uncovering the mechanism of trade-off resolution with phenome-wide association analysis

To elucidate how group composition influences the exploration–vigilance trade-off, we conducted a comprehensive analysis of the phenotypic basis underlying the trade-off alleviation, using a framework we termed phenome-wide higher-level association study (PheHAS). Specifically, we screened 190 phenotypes obtained from the DGRPool database (26) to identify those for which the phenotypic differences between strains were strongly associated with the degree of trade-off alleviation observed across strain pairs. Measures of variability (e.g., SE, SD, or CV) were excluded from analysis.

The trade-off alleviation score for a given strain pair was calculated as follows:

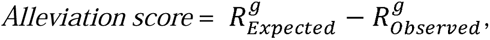

where 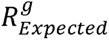 represents the rank expected from the average phenotypic values of the homogeneous groups constituting the focal pair (calculated using Pareto optimization as described above), and 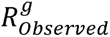 denotes the actual rank of the corresponding heterogeneous group under the same trade-off.

For each strain pair, the phenotypic distance was defined as the absolute difference in a given phenotype between the two strains, across all 190 phenotypes. We computed a distance matrix based on pairwise correlations among the phenotypes and applied hierarchical clustering (Ward’s method), classifying them into 16 clusters. These clusters were further grouped into four higher-order categories based on biological relevance.

For each phenotype, we calculated Pearson’s correlation coefficient between the alleviation score and phenotypic distance across 65 strain pairs, following the method described in our previous study (22). We then applied Fisher’s z-transformation and used a random-effects model to integrate the correlation coefficients within each cluster, yielding pooled estimates and confidence intervals in a meta-analytic framework.

### Statistical analyses

All statistical analyses were conducted using R version 4.3.1. Wilcoxon’s signed rank tests were performed using the ‘ggpubr’ package to evaluate differences in mean values between conditions. Pearson’s correlation analysis was performed to evaluate the association between two variables. The ‘ggeffects’ package was used to predict the mean phenotypic value of each strain or strain pair by considering random effects. We used the ‘rPref’ package to derive relative ranks along the behavioral trade-off dimension, applying Pareto optimization to two indicators: exploration and vigilance (67). Throughout all statistical analyses, standard errors were calculated to represent variation.

## Acknowledgments

We thank Masashi Murakami for his valuable comments on the interpretation and discussion of the results. We are also grateful to the mechanical unit at our university for their kind assistance in constructing the experimental arenas.

## Author contributions

Conceptualization: T.O., D.X.S., Y.T.; Data curation: T.O., D.X.S.; Formal analysis: T.O.; Funding acquisition: Y.T.; Investigation: T.O.; Methodology: T.O., D.X.S.; Project administration: Y.T.; Supervision: D.X.S., Y.T.; Validation: D.X.S., Y.T.; Visualization: T.O.; Writing – original draft: T.O.; Writing – review & editing: T.O., D.X.S., Y.T.

## Data availability

******

## Supplemental figures

**Figure S1.**
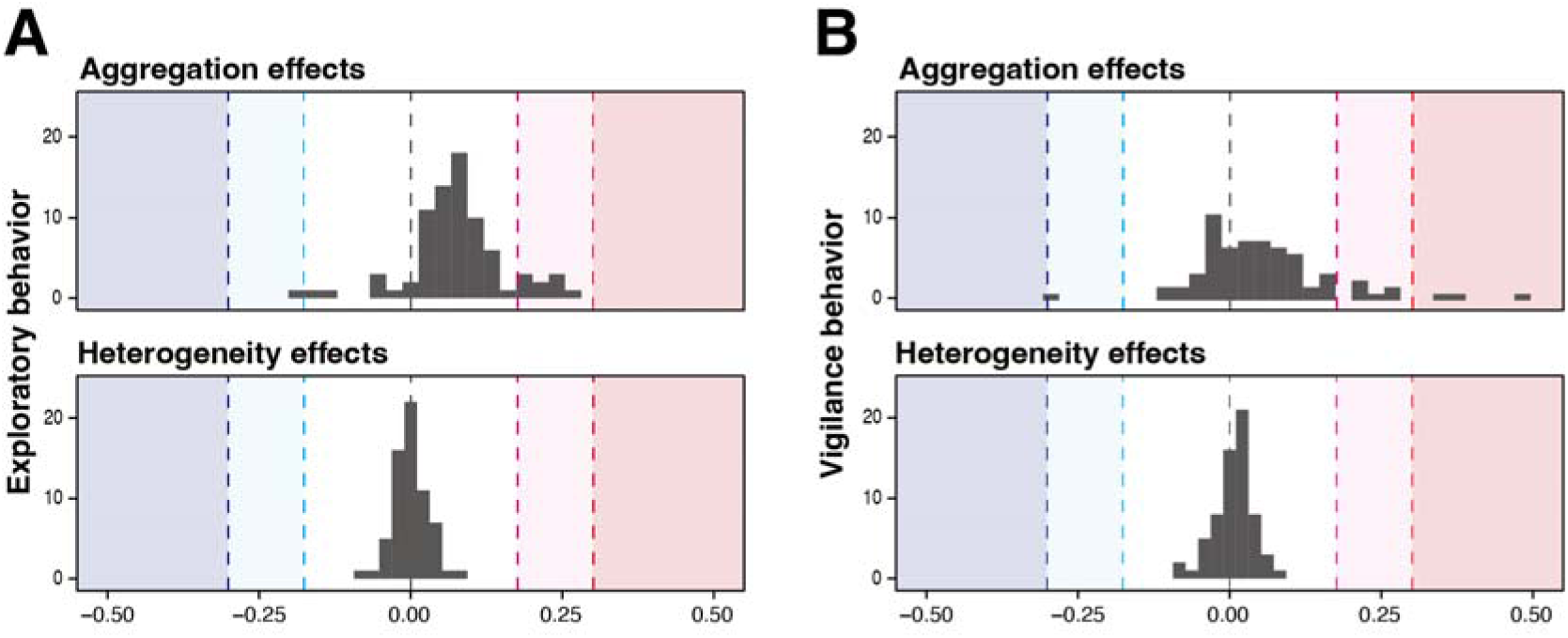
Distribution of aggregation and heterogeneity effects across strains and strain pairs. Histograms show effect size distributions for aggregation (top) and heterogeneity (bottom) in **(A)** exploratory and **(B)** vigilance behavior. Shaded regions mark effect magnitude: red, >2-fold increase; pink, 1.5–2-fold increase; blue, >2-fold decrease; light blue, 1.5– 2-fold decrease. Although overall tendencies were positive (Fig. 2), effect sizes varied widely across strains and strain pairs, emphasizing the importance of social context in group composition.

**Figure S2.**
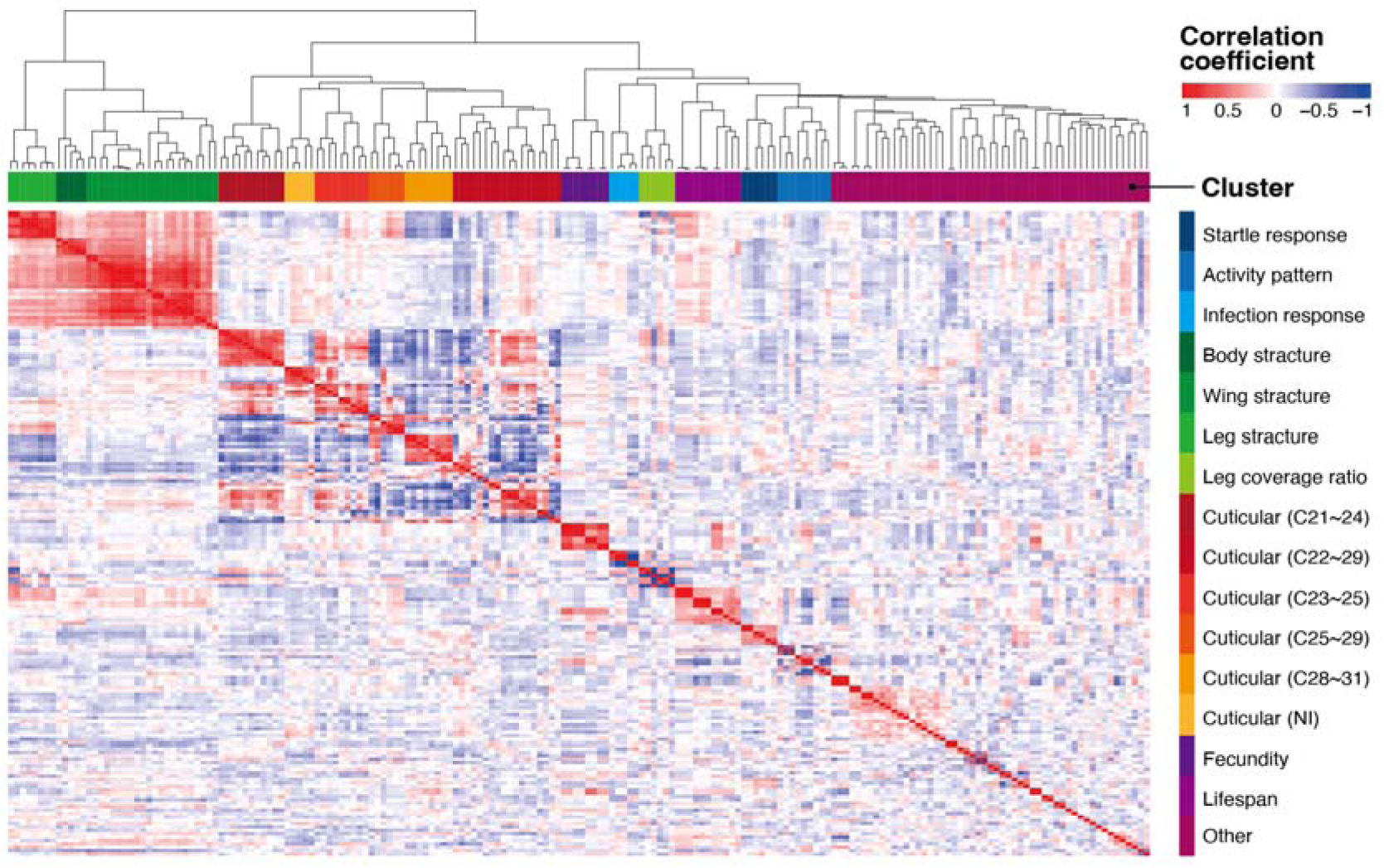
Phenotypic clustering of DGRP strains used for Phenom-wide higher-level association study (PheHAS). Hierarchical clustering of 190 phenotypes from the DGRPool database was performed based on pairwise correlation coefficients (heatmap). The analysis identified 16 clusters, which were further grouped into four higher-order categories: physiology, morphology, cuticular component, and life history. These clusters provided the framework for the phenome-wide higher-level association study (PheHAS), which evaluates how within-group phenotypic variation relates to alleviation of the exploration–vigilance trade-off.

## References

1. S. C. Stearns, Trade-offs in life-history evolution. Funct. Ecol. 3, 259–268 (1989).

2. T. Garland, Trade-offs. Curr. Biol. 24, R60–R61 (2014).

3. M. J. Angilletta, R. S. Wilson, C. A. Navas, R. S. James, Trade-offs and the evolution of thermal reaction norms. Trends Ecol. Evol. 18, 234–240 (2003).

4. J. M. McNamara, A. I. Houston, State-dependent life histories. Nature 380, 215–221 (1996).

5. E. R. Pianka, On r- and K-selection. Am. Nat. 104, 592–597 (1970).

6. W. W. Murdoch, Aspects of the population dynamics of some marsh Carabidae. J. Anim. Ecol. 35, 127–156 (1966).

7. W. W. Murdoch, Population stability and life history phenomena. Am. Nat. 100, 5–11 (1966).

8. J. N. M. Smith, Does high fecundity reduce survival in song sparrows? Evolution 35, 1142–1148 (1981).

9. Th. S. van Dijk, On the relationship between reproduction, age and survival in two carbid beetles. Oecologia 40, 63–80 (1979).

10. R. B. Weladji, et al., Heterogeneity in individual quality overrides costs of reproduction in female reindeer. Oecologia 156, 237–247 (2008).

11. K. A. Hammond, J. Diamond, Maximal sustained energy budgets in humans and animals. Nature 386, 457–462 (1997).

12. A. M. Leroi, A. K. Chippindale, M. R. Rose, Long-term laboratory evolution of a genetic life-history trade-off in *Drosophila melanogaster*. 1. The role of genotype-by-environment interaction. Evolution 48, 1244–1257 (1994).

13. D. Reznick, L. Nunney, A. Tessier, Big houses, big cars, superfleas and the costs of reproduction. Trends Ecol. Evol. 15, 421–425 (2000).

14. J. S. Brown, Vigilance, patch use and habitat selection: Foraging under predation risk. Evol. Ecol. Res. 1, 49–71 (1999).

15. S. Lima, L. Dill, Behavioral decisions made under the risk of predation: A review and prospectus. Can. J. Zool. 68, 619–640 (1990).

16. J. Balaban-Feld, et al., Individual willingness to leave a safe refuge and the trade-off between food and safety: A test with social fish. Proc. R. Soc. B Biol. Sci. 286, 20190826 (2019).

17. B. C. R. Bertram, Vigilance and group size in ostriches. Anim. Behav. 28, 278–286 (1980).

18. G. Beauchamp, Group-size effects on vigilance: A search for mechanisms. Behav. Processes 63, 111–121 (2003).

19. G. Beauchamp, What is the magnitude of the group-size effect on vigilance? Behav. Ecol. 19, 1361–1368 (2008).

20. J. W. Jolles, A. J. King, S. S. Killen, The role of individual heterogeneity in collective animal behaviour. Trends Ecol. Evol. 35, 278–291 (2020).

21. M. Nagy, et al., Synergistic benefits of group search in rats. Curr. Biol. 30, 4733–4738.e4 (2020).

22. T. Okuyama, D. X. Sato, Y. Takahashi, Genetic heterogeneity induces non-additive behavioural changes in *Drosophila*. J. Exp. Biol. 228, jeb249449 (2025).

23. F. Santostefano, A. J. Wilson, P. T. Niemelä, N. J. Dingemanse, Indirect genetic effects: A key component of the genetic architecture of behaviour. Sci. Rep. 7, 10235 (2017).

24. E. W. Wice, J. B. Saltz, Indirect genetic effects for social network structure in *Drosophila melanogaster*. Philos. Trans. R. Soc. B Biol. Sci. 378, 20220075 (2023).

25. T. F. C. Mackay, et al., The *Drosophila melanogaster* genetic reference panel. Nature 482, 173–178 (2012).

26. V. Gardeux, et al., DGRPool: A web tool leveraging harmonized *Drosophila* Genetic Reference Panel phenotyping data for the study of complex traits. eLife 12 (2024).

27. D. X. Sato, Y. Takahashi, Neurogenomic and behavioral principles shape freezing dynamics and synergistic performance in *Drosophila melanogaster*. Nat. Commun. 16, 5928 (2025).

28. W. D. Hamilton, Geometry for the selfish herd. J. Theor. Biol. 31, 295–311 (1971).

29. S. R. J. Neill, J. M. Cullen, Experiments on whether schooling by their prey affects the hunting behaviour of cephalopods and fish predators. J. Zool. 172, 549–569 (1974).

30. A. A. Fonseca, C. A. dos Santos, Reviewing social facilitation in insects over the past 30 years. Sociobiology 70, e9210 (2023).

31. R. B. Zajonc, Social facilitation. Science 149, 269–274 (1965).

32. K. Connolly, The social facilitation of preening behaviour in *Drosophila melanogaster*. Anim. Behav. 16, 385–391 (1968).

33. A. Muria, et al., Social facilitation of long-lasting memory is mediated by CO_2_ in *Drosophila*. Curr. Biol. 31, 2065–2074.e5 (2021).

34. M.-A. Chabaud, G. Isabel, L. Kaiser, T. Preat, Social facilitation of long-lasting memory retrieval in *Drosophila*. Curr. Biol. 19, 1654–1659 (2009).

35. K. Roelofs, Freeze for action: Neurobiological mechanisms in animal and human freezing. Philos. Trans. R. Soc. B Biol. Sci. 372, 20160206 (2017).

36. R. J. Egan, et al., Understanding behavioral and physiological phenotypes of stress and anxiety in zebrafish. Behav. Brain Res. 205, 38–44 (2009).

37. C. E. Howard, et al., Serotonergic modulation of walking in *Drosophila*. Curr. Biol. 29, 4218–4230.e8 (2019).

38. S. L. Lima, Back to the basics of anti-predatory vigilance: The group-size effect. Anim. Behav. 49, 11–20 (1995).

39. J. E. Treherne, W. A. Foster, Group transmission of predator avoidance behaviour in a marine insect: The trafalgar effect. Anim. Behav. 29, 911–917 (1981).

40. T. H. Clutton-Brock, et al., Selfish sentinels in cooperative mammals. Science 284, 1640–1644 (1999).

41. P. A. Bednekoff, Mutualism among safe, selfish sentinels: A dynamic game. Am. Nat. 150, 373–392 (1997).

42. B. K. Paul, et al., Crop-livestock integration provides opportunities to mitigate environmental trade-offs in transitioning smallholder agricultural systems of the Greater Mekong Subregion. Agric. Syst. 195, 103285 (2022).

43. K. Ohashi, A. Jürgens, J. D. Thomson, Trade-off mitigation: A conceptual framework for understanding floral adaptation in multispecies interactions. Biol. Rev. 96, 2258–2280 (2021).

44. S. N. Beshers, J. H. Fewell, Models of division of labor in social insects. Annu. Rev. Entomol. 46, 413–440 (2001).

45. E. Bonabeau, G. Theraulaz, J.-L. Deneubourg, Quantitative study of the fixed threshold model for the regulation of division of labour in insect societies. Proc. Biol. Sci. 263, 1565–1569 (1996).

46. U. Klibaite, J. W. Shaevitz, Paired fruit flies synchronize behavior: Uncovering social interactions in *Drosophila melanogaster*. PLOS Comput. Biol. 16, e1008230 (2020).

47. P. Ramdya, et al., Mechanosensory interactions drive collective behaviour in *Drosophila*. Nature 519, 233–236 (2015).

48. S. Sarin, R. Dukas, Social learning about egg-laying substrates in fruitflies. Proc. R. Soc. B Biol. Sci. 276, 4323–4328 (2009).

49. E. Danchin, et al., Cultural flies: Conformist social learning in fruitflies predicts long-lasting mate-choice traditions. Science 362, 1025–1030 (2018).

50. R. B. Rosengaus, A. B. Maxmen, L. E. Coates, J. F. A. Traniello, Disease resistance: A benefit of sociality in the dampwood termite *Zootermopsis angusticollis* (Isoptera: Termopsidae). Behav. Ecol. Sociobiol. 44, 125–134 (1998).

51. S. Cremer, S. A. O. Armitage, P. Schmid-Hempel, Social immunity. Curr. Biol. 17, R693–R702 (2007).

52. A. Fraimout, et al., Phenotypic plasticity of *Drosophila suzukii* wing to developmental temperature: Implications for flight. J. Exp. Biol. 221, jeb166868 (2018).

53. V. Godesberg, T. Bockemühl, A. Büschges, Natural variability and individuality of walking behavior in *Drosophila*. J. Exp. Biol. 227, jeb247878 (2024).

54. H. Dankert, L. Wang, E. D. Hoopfer, D. J. Anderson, P. Perona, Automated monitoring and analysis of social behavior in *Drosophila*. Nat. Methods 6, 297–303 (2009).

55. J. M. Mueller, N. Zhang, J. M. Carlson, J. H. Simpson, Variation and variability in *Drosophila* grooming behavior. Front. Behav. Neurosci. 15, 769372 (2022).

56. H. Chung, S. B. Carroll, Wax, sex and the origin of species: Dual roles of insect cuticular hydrocarbons in adaptation and mating. BioEssays 37, 822–830 (2015).

57. R. W. Howard, G. J. Blomquist, Ecological, behavioral, and biochemical aspects of insect hydrocarbons. Annu. Rev. Entomol. 50, 371–393 (2005).

58. M. H. Gromko, T. A. Markow, Courtship and remating in field populations of *Drosophila*. Anim. Behav. 45, 253–262 (1993).

59. G. Arnqvist, T. Nilsson, The evolution of polyandry: Multiple mating and female fitness in insects. Anim. Behav. 60, 145–164 (2000).

60. T. Chapman, L. F. Liddle, J. M. Kalb, M. F. Wolfner, L. Partridge, Cost of mating in *Drosophila melanogaster* females is mediated by male accessory gland products. Nature 373, 241–244 (1995).

61. D. X. Sato, T. Okuyama, Y. Takahashi, Multifaceted and extensive behavioral trajectories of genomically diverse *Drosophila* lines. Sci. Data 12, 400 (2025).

62. R. Zacarias, S. Namiki, G. M. Card, M. L. Vasconcelos, M. A. Moita, Speed dependent descending control of freezing behavior in *Drosophila melanogaster*. Nat. Commun. 9, 3697 (2018).

63. C. H. Ferreira, M. A. Moita, Behavioral and neuronal underpinnings of safety in numbers in fruit flies. Nat. Commun. 11, 4182 (2020).

64. E. Eyjolfsdottir, et al., Detecting social actions of fruit flies in In Computer Vision – ECCV 2014, D. Fleet, T. Pajdla, B. Schiele, T. Tuytelaars, Eds. (Springer International Publishing, 2014), pp. 772–787.

65. E. H. Simpson, Measurement of diversity. Nature 163, 688–688 (1949).

66. S. Borzsony, D. Kossmann, K. Stocker, The Skyline operator. in Proceedings 17th International Conference on Data Engineering, (2001), pp. 421–430.

67. P. Roocks, Computing Pareto frontiers and database preferences with the rPref package. R J. 8, 393–404 (2016).

